# Na^+^/Ca^2+^ exchangers and Orai channels jointly refill endoplasmic reticulum (ER) Ca^2+^ via ER nanojunctions in vascular endothelial cells

**DOI:** 10.1101/084285

**Authors:** Cristiana M. L. Di Giuro, Niroj Shrestha, Roland Malli, Klaus Groschner, Cornelis van Breemen, Nicola Fameli

## Abstract

We investigated the role of Na^+^/Ca^2+^ exchange (NCX) in the refilling of endoplasmic reticulum (ER) Ca^2+^ in vascular endothelial cells under various conditions of cell stimulation and plasma membrane (PM) polarization. Better understanding of the mechanisms behind basic ER Ca^2+^ content regulation is important, since current hypotheses on the possible ultimate causes of ER stress point to deterioration of the Ca^2+^ transport mechanism to/from ER itself. We measured [Ca^2+^]_i_ temporal changes by Fura-2 fluorescence under experimental protocols that inhibit a host of transporters (NCX, Orai, non-selective transient receptor potential canonical (TRPC) channels, sarco/endoplasmic reticulum Ca^2+^ ATPase (SERCA), Na^+^/K^+^ ATPase (NKA)) involved in the Ca^2+^ communication between the extracellular space and the ER. Following histamine-stimulated ER Ca^2+^ release, blockade of NCX Ca^2+^-influx mode (by 10 *µ*M KB-R7943) diminished the ER refilling capacity by about 40%, while in Orai1 dominant negative-transfected cells NCX blockade attenuated ER refilling by about 60%. Conversely, inhibiting the high-ouabain-affinity NKA (10 nM ouabain), which may be localized in PM-ER junctions, increased the ER Ca^2+^ releasable fraction by about 20%, thereby supporting the hypothesis that this process of privileged ER refilling is junction-mediated. Junctions were observed in the cell ultrastructure and their main parameters of membrane separation and linear extension were (9.6 *±* 3.8) nm and (128 *±* 63) nm, respectively. Our findings point to a process of privileged refilling of the ER, in which NCX and SOCE via the stromal interaction molecule (STIM)-Orai system are the sole protagonists. These results shed light on the molecular machinery involved in the function of a previously hypothesized subplasmalemmal Ca^2+^ control unit during ER refilling with extracellular Ca^2+^.

## 1 Introduction

Activation of non-excitable cells, like endothelial cells (EC), by receptor agonists typically provokes an increase in inositol 1,4,5-trisphosphate (IP_3_) and consequent endoplasmic reticulum (ER) Ca^2+^ release via IP_3_ receptors (IP_3_R). The subsequent [Ca^2+^]_i_ elevation is a preamble underpinning basic EC function like nitric oxide (NO) synthesis by the endothelial NO synthase and endothelin-1 production [1–4]. While store-operated Ca^2+^ entry (SOCE) is generally considered the main (if not sole) mechanism responsible for cytosolic Ca^2+^ resetting and ER Ca^2+^ refilling following stimulation [5, 6], and evidence exists for a role of mitochondria in ER Ca^2+^ regulation [5, 7], there is reason to consider that Na^+^/Ca^2+^ exchange (NCX) may also play an important role in this process. In fact, the Na^+^*/*Ca^2+^ exchanger (more specifically we will refer herein to the NCX1 isoform—the most abundantly expressed in EC [8, 9]) has been shown to play an important role in Ca^2+^ extrusion from these cells once the stimulus is removed in low external Ca^2+^ conditions as well as during acetylcholine and histamine stimulation under physiological conditions [10–12]. In addition to its Ca^2+^ extrusion operation, the NCX Ca^2+^ influx mode has been observed to perform a fundamental function in Ca^2+^ refilling of sarco/endoplasmic reticulum (S/ER) in vascular smooth muscle cells [13] and it has also been suggested that it plays a role in activating angiogenesis function in EC [14]. Therefore, since the presence of NCX in endothelial cells has been reported [15], we studied in depth the role of NCX for ER Ca^2+^ refilling in human umbilical vein derived EC (EA.hy926).

An important feature of NCX-mediated S/ER refilling in earlier observations, in vascular smooth muscle cells as well as EC, is that it appears to occur via plasma membrane (PM)-S/ER nanojunctions [10, 13, 16, 17], intracellular regions comprising portions of the PM and of the S/ER membrane separated by 10–30 nm and extending roughly parallel to each other for several hundred nm, the intervening cytosolic space and the local ion transport machinery (channels, pumps, exchangers). The role of nanojunctions in assisting cell signaling has been recognized as pivotal in several systems [18, 19]. Interestingly, there is also strong evidence that the subplasmalemmal region of EC is involved in a crucial way in Ca^2+^ extrusion mechanisms, in a manner that shunts the bulk cytoplasm; indeed, this has led to the hypothesis of a “subplasmalemmal Ca^2+^ control unit (SCCU)” akin to the “superficial buffer barrier” of vascular smooth muscle cells [20–22]. Hence, in the present study we also addressed the important hypothesis that NCX-mediated Ca^2+^ transport in EC occurs via PM-ER junctions.

To provide the right setting of our findings, it is useful to recall the current knowledge of the machinery purportedly responsible for Ca^2+^ handling in endothelial cells. Due to the apparent lack of a functional role for voltage-gated cation channels in EC [23, 24], Ca^2+^ entry in these cells depends upon receptor operated channels, predominantly channels of the non-selective transient receptor potential canonical (TRPC) family, and the highly Ca^2+^ selective Orai1 store operated channels, the latter only operative when clustered in PM-ER junctions, upon depletion of the ER Ca^2+^ [25]. Furthermore, Ca^2+^ can be pumped into the ER by the S/ER Ca^2+^ ATPases (SERCA), and released from the ER via the IP_3_R and ryanodine receptor channels. However, while the latter have been observed by immunofluorescence [26] and implicated in the ER Ca^2+^ release in the subplasmalemmal region of EC [10–12, 20, 21], they do not appear to be expressed in the human umbilical vein derived endothelial cell line employed in this project (EA.hy926) [27].

Due to the reported variability of the EC’s membrane potential, *V*_m_, (see [28] and references therein) and the necessity to operate with a stable driving force of the NCX, we opted to work under conditions of membrane depolarization by using a superfusing solution with 70 mM [K^+^] (with *≈* 70 mM [Na^+^] for osmotic balance). This “driving force clamp” method has been employed before [29] and in our experiments also serves the purpose of highlighting the NCX-mediated Ca^2+^ influx in relation to that of TRPC and Orai1.

In the present study, we report Fura-2-based [Ca^2+^]_i_ measurements under a host of conditions aimed at elucidating the role of NCX in the refilling of ER Ca^2+^ via PM-ER nanojunctions. We also provide novel ultrastructural evidence of PM-ER junctions and quantitative characterization thereof in EC.

## 2 Materials and methods

### 2.1 Materials

Cell culture chemicals were obtained from Thermo Fisher Scientific; histamine, amphotericin B (AmB), KCl, MgCl_2_, CaCl_2_, glucose, HEPES, EGTA, 2-Aminoethoxydiphenyl borate (2-APB) and ouabain from Sigma-Aldrich, NaCl from Carl Roth. KB-R7943 was purchased from Sigma-Aldrich and Abcam. We used Fura-2 AM products by both Sigma-Aldrich and Thermo Fisher Scientific.

### 2.2 Cell culture and transfection

Experiments were performed with the human umbilical vein endothelium derived cell line, EA.hy926. Cells were cultured in Dulbecco’s Modified Eagle Medium (DMEM) containing 10% fetal bovine serum, 100 U/mL penicillin, 100 *µ*g/mL streptomycin and HAT (50 *µ*M hypoxanthin, 0.2 *µ*M aminopterin, 0.8 *µ*M thymidine and 1.25 *µ*g/mL AmB) and were maintained in an incubator at 37^*°*^C in 5% CO_2_ atmosphere. The cells were used at passages between 30 and 80 and were plated on 6 mm *×* 6 mm square cover slips for transfection and experiments at a confluence of 50–80%.

Cells were transiently transfected with 1.5 *µ*g Orai1^dn^ cDNA using TransFast^TM^ Transfection Reagent (Promega) in DMEM following manufacturers instructions. After 4–6 h incubation the transfection mixture was replaced by normal culture medium. Ca^2+^ measurements were performed 24–48 h after transfection.

### 2.3 Solutions

The normal experimental buffer (EB) was composed of (in mM): 138 NaCl, 5 KCl, 2 CaCl_2_, 1 MgCl_2_, 10 glucose and 10 HEPES, pH adjusted to 7.4 with NaOH.

The 70 mM [K^+^]_o_ EB was composed of (in mM): 73 NaCl, 70 KCl, 2 CaCl_2_, 1 MgCl_2_, 10 glucose and 8 HEPES, pH adjusted to 7.4 with KOH. The Ca^2+^-free EB contained 0.1 mM EGTA instead of CaCl_2_.

The loading buffer solution (LB) contained (in mM unless otherwise indicated): 135 NaCl, 5 KCl, 2 CaCl_2_, 1 MgCl_2_, 10 HEPES, 2.6 NaHCO_3_, 0.44 KH_2_PO_4_, 10 glucose with 0.1% vitamins and 0.2% essential amino acids, 1% penicillin/streptomycin, 1.25 *µ*g/mL AmB, pH adjusted to 7.4 with NaOH.

### 2.4 Single-cell cytosolic [Ca^2+^]_i_ measurements

Changes in [Ca^2+^]_i_ were monitored using the Fura-2 technique as previously described [30]. Briefly, cells on cover slips were loaded with 2 *µ*M Fura-2 AM for 45 min in LB at room temperature in the dark. After the incubation period, cells were washed twice with LB, and left to equilibrate for at least 20 min also in LB. The coverslip was then mounted in a perfusion chamber on an inverted microscope (Olympus IX71) and perfused with different solutions at room temperature. During the recordings using Live Acquisition 2.5 software (FEI, Germany), cells were excited alternately at 340 and 380 nm using an Oligochrome excitation system (FEI, Germany) and fluorescent images were captured at 510 nm every 1 s with an ORCA-03G digital CCD camera (Hamamatsu, Germany).

### 2.5 Electron microscopy

EA.hy926 cells were grown on an Aclar film substrate (Gr¨opl, Tulln, Austria). The primary fixative solution contained 2.5% glutaraldehyde and 2% paraformaldehyde (Ted Pella, Redding, CA, USA) in 0.1 M phosphate buffer, pH 7.4, for 45 min.

In the process of secondary fixation, the cell sets were fixed with either 2% OsO_4_ or 1% OsO_4_ + 1% K_3_Fe(CN)_6_ for 45 min at room temperature. The samples were then dehydrated in increasing concentrations of acetone (25, 50, 70, 80, 90, and 95%) and in the final process of dehydration, the samples underwent 3 washes in 100% acetone. The cells were then resin-infiltrated in increasing concentrations of TAAB resin (30, 50, and 70% in acetone). The infiltration process was completed by three passages in 100% resin. The cells were finally resin-embedded in molds and polymerized at 60^*°*^C for 3 days.

To produce the imaging samples, 80-nm sections were cut from the embedded sample blocks with a UC 7 Ultramicrotome (Leica Microsystems, Vienna, Austria) using a diamond knife (Diatome, Biel, Switzerland) and were collected on Pioloform-coated 100-, 200- and 300-mesh copper grids (hexagonal or square meshes; Gr¨opl, Tulln, Austria). The sections were post-stained with Pt-blue and Reynolds lead citrate for 15 and 7 minutes, respectively.

Electron micrographs at various magnifications were obtained with a Tecnai G 2 FEI transmission electron microscope at 120 kV and equipped with an ultrascan 1000cd camera (Gatan).

Cells were imaged both in a “plan” view, i.e., looking down perpendicular to the plane of the substrate on which the cells were grown, and in an “elevation” orientation, that is, observing the cells along a direction parallel to the plane of their substrate. This was done in an effort to capture as many instances of PM-ER junctions as possible and get around the difficulty of sectioning the PM effectively in the plan view due to the flatness and thinness of the cells especially at their periphery. Elevation-view blocks were prepared by cutting out a portion of a plan-view block and re-embedding it in resin tilted through 90 degrees with respect to its original orientation.

### 2.6 Data analysis, statistics, software

Individual experiments in each reported set of *n* consist of Fura-2 signal trace measurements from approximately 5 to 20 cells. Means, standard deviations (SD) and standard errors (SEM) are then calculated for each experiment. After a set of experiments, the pooled statistics are, in turn, calculated. The reported traces are the values of the means *±* pooled SD. In our statistical tests, *p*-values *≤* 0.05 were considered significant.

Given the slightly different behaviour observed in different sub-populations of cells, our approach to determining significance in the effects recorded under the stimuli and inhibitors used in our experiments was to analyze the ratio between the amplitudes of the Ca^2+^ transients generated by the various conditions as a measure of ER Ca^2+^ content and refilling efficiency and of cytosolic [Ca^2+^] changes. Therefore, after calculating the mean Ca^2+^ signal traces from each type of experiment and referring to the protocol description in section 3.1, we report our data as the amplitude ratios of the 2nd histamine transient to the 1st and of the nCa^2+^ phase transient to the 2nd histamine one. A third interesting parameter for our analysis is the rate of Ca^2+^ re-entry in the cytosol during the nCa^2+^ phase. This is calculated as the slope of the roughly linear portion of the Fura-2 signal at the onset of the nCa^2+^ phase.

The Fura-2 340/380 fluorescence ratios were recorded as text files and then analyzed by a combination of in-house C programs, Linux shell scripts and LibreOffice Calc Spreadsheet functions (http://www.libreoffice.org).

The electron micrographs were recorded as TIFF image files and analyzed with xfig (http://www.xfig.org) to measure PM-ER junctional separations and extensions. Plots were generated with gnuplot (http://gnuplot.info).

We used JabRef as reference manager [31].

## 3 Results

Present knowledge suggests that the essential Ca^2+^ handling machinery at work in EC comprises TRPC and Orai1 channels, NCX and the PM Ca^2+^ ATPase (PMCA) on the PM, SERCA, and IP_3_R on the ER. Less directly involved with Ca^2+^ handling, are the NKA pumps on the PM [32]. The schematic diagram in figure 1 provides a picture of how these players might be implicated in the shaping of Ca^2+^ signals in EC. Starting from the hypothesis that Ca^2+^ entry via the NCX is responsible at least in part for ER Ca^2+^ refilling and that this NCX-mediated refilling takes place via PM-ER junctions, we elucidated the role of the NCX by systematically inhibiting or eliminating all the other potential sources of Ca^2+^ entry in EC by the employment of pharmacological agents and gene transfer techniques, in a manner that would also highlight the possible involvement of junctional transport.

### 3.1 ER Ca^2+^ is refilled by extracellular Ca^2+^ regardless of membrane polarization

Our main experimental protocol and basic results are displayed in figure 2. The cells were kept under normal physiological conditions for the first min, then superfused with nominally Ca^2+^ free (0Ca^2+^) buffer for 3 min, after which they were bathed for a further 3 min in 0Ca^2+^ buffer with 100 *µ*M histamine, to initiate ER Ca^2+^ release—by increasing IP_3_ production and, in turn, sensitizing IP_3_R to Ca^2+^—noticeable as a Ca^2+^ transient during this phase of the protocol (the 0Ca^2+^ buffer choice ensures that the observed transient at this stage is solely due to ER Ca^2+^ mobilization). Histamine is then eliminated by a 3-min washout with 0Ca^2+^ buffer, before superfusing cells with normal Ca^2+^ (nCa^2+^) buffer for 4 min again to allow extracellular Ca^2+^ influx after ER Ca^2+^ release. Upon completion of the Ca^2+^ influx phase, the cells are returned to 0Ca^2+^ buffer before administering a second 3-min histamine stimulus aimed at revealing the amount of ER refilling that took place during the nCa^2+^ phase of the protocol. Clearly, the extracellular Ca^2+^ influx taking place during this phase manifests itself in a global [Ca^2+^]_i_ rise (central transient in figure 2B) and is able to refill the ER of the majority of its original content, as revealed by the second ER Ca^2+^ release amounting to about 67% of the first (figure 2B and D).

**Figure 1:**
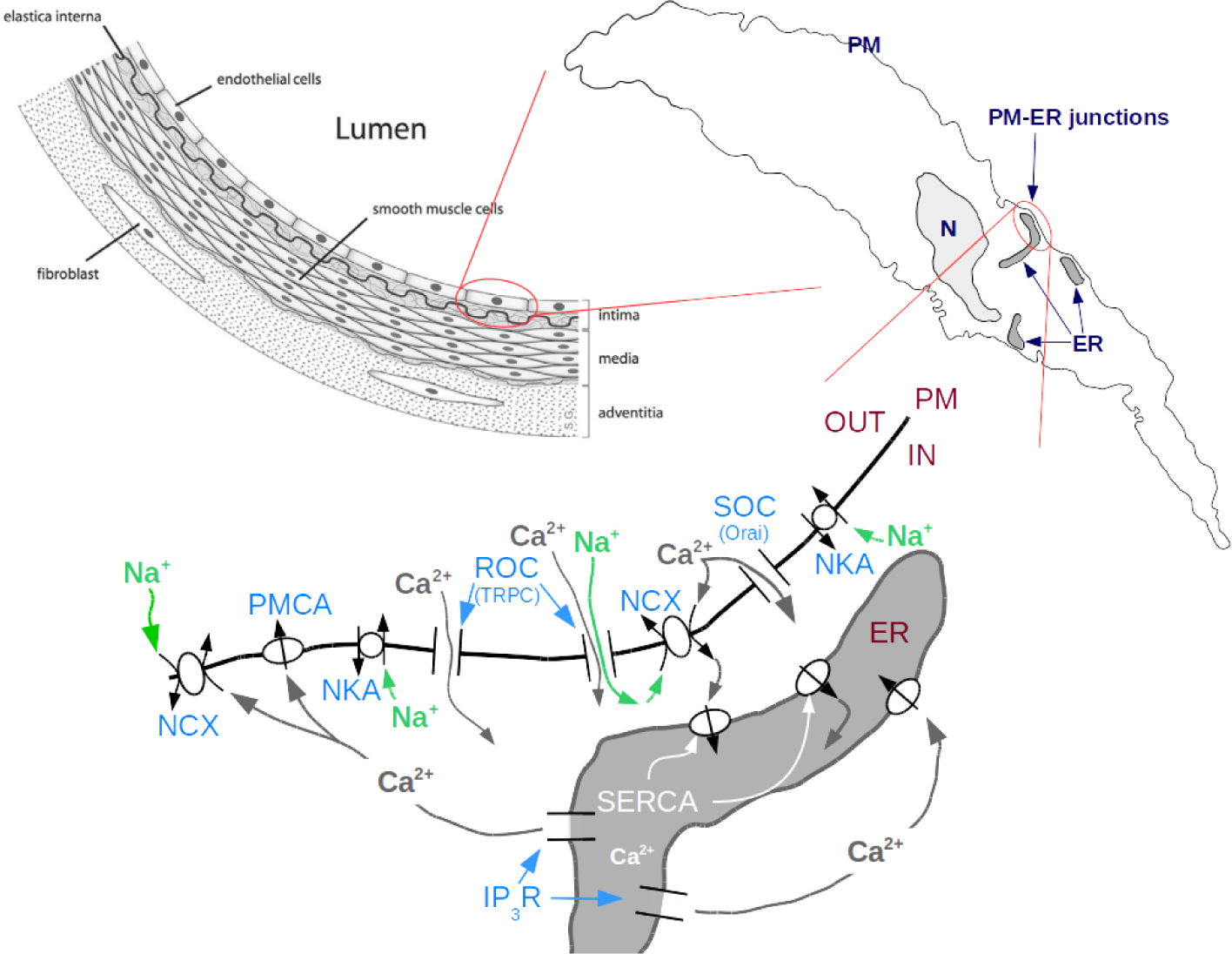
Top left: schematic partial cross-section of a vessel with location of EC (image attribution: DRosenbach at English Wikipedia); top right: diagram of one EC showing location of PM-ER junctions; bottom: essential depiction of the current understanding of the Ca^2+^ transport machinery and some of the mechanisms responsible for ER Ca^2+^ refilling in EC. (N = nucleus; ROC = receptor operated channels; SOC = store operated channels; see text for the other acronyms.)

Since one of the hypotheses of the study is to understand the role of the Ca^2+^ influx mode of the NCX in the ER Ca^2+^ refilling mechanism, in all of our subsequent experiments, from 2 min prior to the nCa^2+^ phase onwards, we superfused the cells in 70 mM [K^+^] buffer to depolarize the cell membrane, as a way to minimize Ca^2+^ entry via TRPC and SOCE via the STIM-Orai1 system, and thereby emphasizing the contribution of the NCX. This situation is depicted in figure 2A.

Moreover, depolarizing the membrane essentially clamps its potential at a value around 0 mV [28], which holds fixed the driving force of the NCX, given by the relationship *E*_NCX_ < *V*_m_ between the NCX reversal potential, *E*_NCX_, and the membrane potential, *V*_m_. This is important since the *V*_m_ of EC (EA.hy926 included) has been shown to be rather variable [28, 33, 34] and this variability will also reflect in the driving force of the NCX.

**Figure 2:**
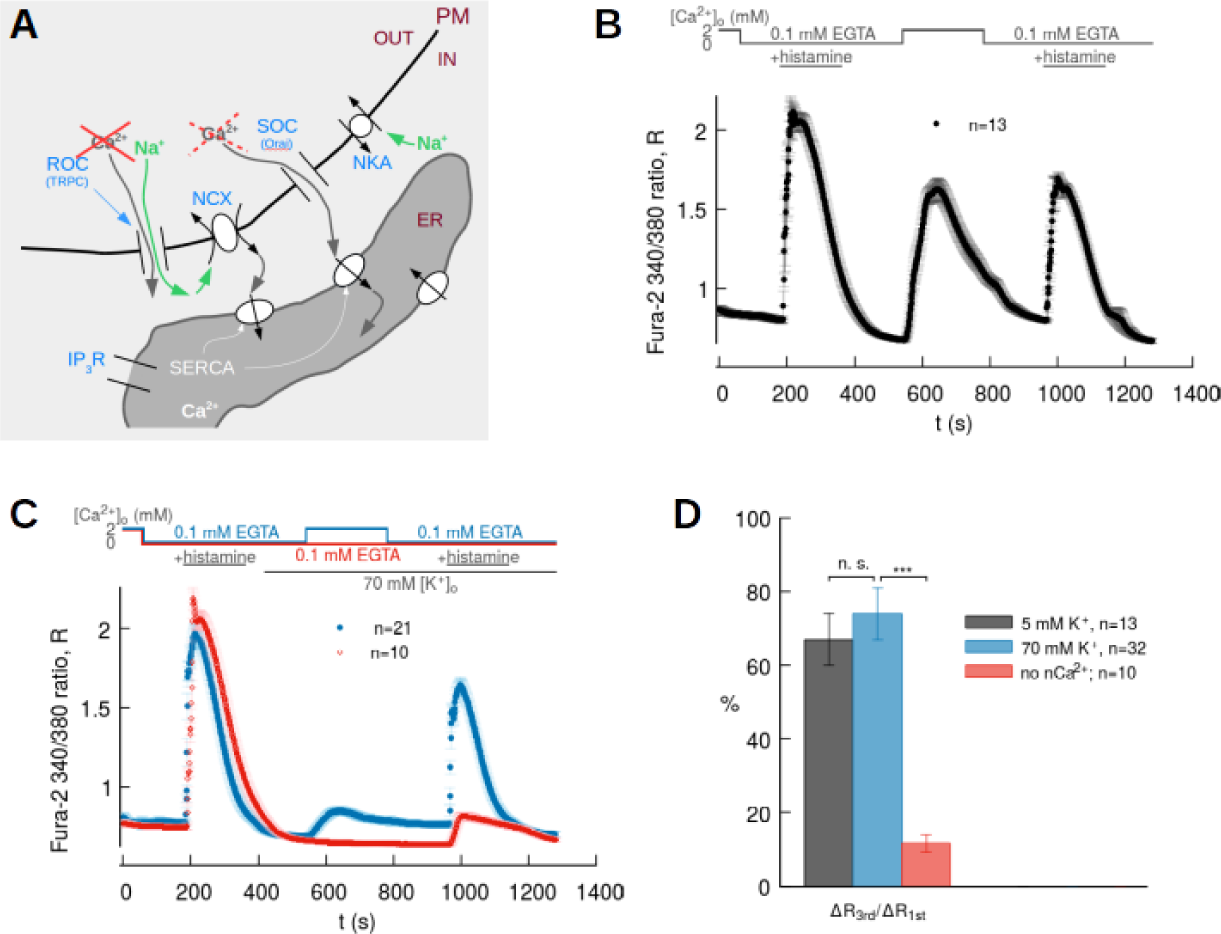
**A**. Schematics of the possible ion fluxes between the extra-cellular space and ER under 70-mM-[K^+^]_o_ depolarizing condition; **B**. Main protocol results under normal polarization conditions: [Ca^2+^]_o_ and histamine stimulation timeline (top), recorded Fura-2 signal (bottom); **C**. Effect of depolarization on Ca^2+^ entry and ER refilling (solid blue circles); effect of nCa^2+^ phase omission on the Ca^2+^ signal (open red diamonds); **D**. Bar chart from traces in B, C.

Under these experimental conditions, we recorded a greatly diminished global [Ca^2+^]_i_ signal during the nCa^2+^ phase, while the ER is refilled to the same level observed in polarized cells (solid blue circles in figure 2C and D).

In this set of experiments, we also determined the contribution of extracellular Ca^2+^ as a source of ER Ca^2+^ refilling by omitting the nCa^2+^ phase from our main protocol; we report these data in red in figure 2C and D. Evidently, when external Ca^2+^ is virtually absent, there is no noticeable Ca^2+^ (re-)entry in the cytoplasm and the ER Ca^2+^ release peak is significantly lower than when extracellular Ca^2+^ is present in the superfusate. We also observed that the remnant ER Ca^2+^ in this case is largely due to re-uptake by SERCA pumps during the first ER Ca^2+^ release, since, by blocking SERCA with 2,5-Di-tert-butyl-1,4-benzoquinone (BHQ) during the first histamine stimulation, we found that the Ca^2+^ left after the second stimulation under 0Ca^2+^ conditions throughout the protocol was even further reduced (data not shown).

### 3.2 NCX contribution to ER Ca^2+^ refilling

To examine whether and to what extent the NCX-mediated Ca^2+^ influx plays a role in ER refilling, we inhibited the NCX Ca^2+^ influx mode by adding 10 *µ*M of KB-R7943 to the superfusing solution in conjunction with the cell depolarization phase of our protocol (figure 3; in this and the following 2 figures the traces obtained in the depolarization-only experiments—solid blue circles—are reported as reference).

Ca^2+^ release following the nCa^2+^ phase of the protocol shows that the ER Ca^2+^ refilling is about 40% less than in the reference case without KB-R7943 administration (figure 3B and 3C left). While the Ca^2+^ re-entry transient amplitude is comparable to the same phase in the reference data (figure 3C right), the average rate of re-entry during the initial nCa^2+^ phase is markedly different in the two cases (figure 3D).

### 3.3 NCX-mediated ER Ca^2+^ refilling takes place via PM-ER junctions

The observations collected to this point provide a strong indication that the ER Ca^2+^ refilling mechanism under membrane depolarization conditions is all but transparent to the bulk signal. If this were an indication that the NCX-mediated refilling is assisted by PM-ER junctions, we should see differences in the Ca^2+^ signal if we somehow interfered with junctional transport.

We attempted this by inhibiting the low-ouabain-affinity pumps NKA*α*_2_ and NKA*α*_3_, which are known in certain cell types, like vascular smooth muscle, to reside primarily in PM-ER junctions [35–37]. As qualitatively depicted in the schematics of figure 4A, due to the increased junctional Na^+^ accumulation, this should enhance Ca^2+^ entry via NCX, increase ER refilling and in essence provoke the opposite effect of KB-R7943 inhibition, that is, a higher ER Ca^2+^ release signal upon 2^nd^ histamine stimulation. Our findings in this set of experiments corroborate the expectations, in that our measured Ca^2+^ transient amplitude during the 2^nd^ histamine-stimulated release indicates that ER Ca^2+^ refilling levels are on average 20% higher than in the experiments without inhibition (figure 4B and 4C left). In those experiments too, we found that the nCa^2+^phase transient amplitude did not differ appreciably from the reference data (4C right), while the rate of entry at the onset of the nCa^2+^ phase was significantly different (4D).

**Figure 3:**
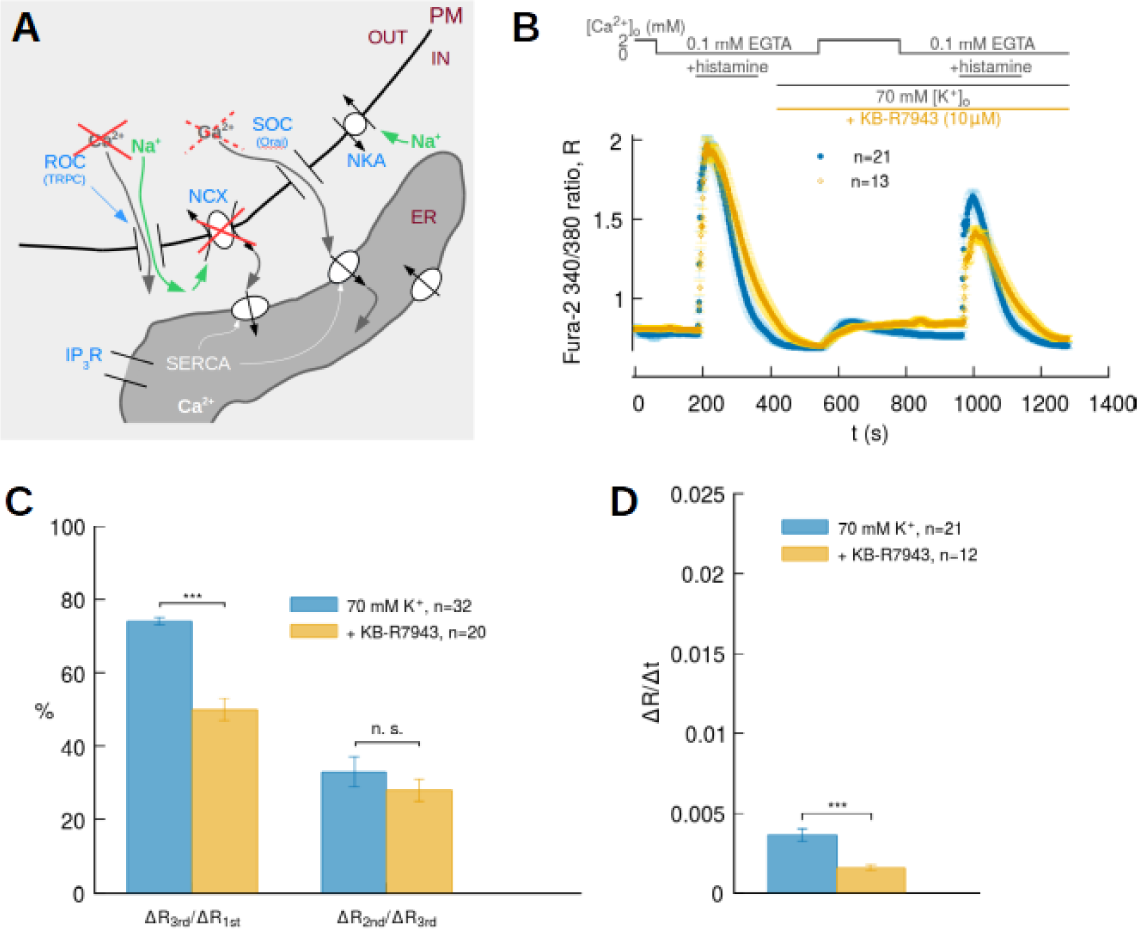
**A**. Schematics of the possible ion fluxes at PM-ER junction locations during the NCX Ca^2+^ influx mode inhibition experiments; **B**. Effect of 10 *µ*M KB-R7943 on ER refilling (open yellow diamonds); recordings from our standard protocol (solid blue circles) reported as reference; **C**. Bar chart from traces in B; left, ratio between 3^rd^ and 1^st^ Ca^2+^ transient amplitudes in A; right, ratio between 2^nd^ and 3^rd^ amplitudes in A; **D**. Bar chart from traces in B, comparing the Ca^2+^ re-entry rates into the cytoplasm at the start of the nCa^2+^ phase of the protocol.

**Figure 4:**
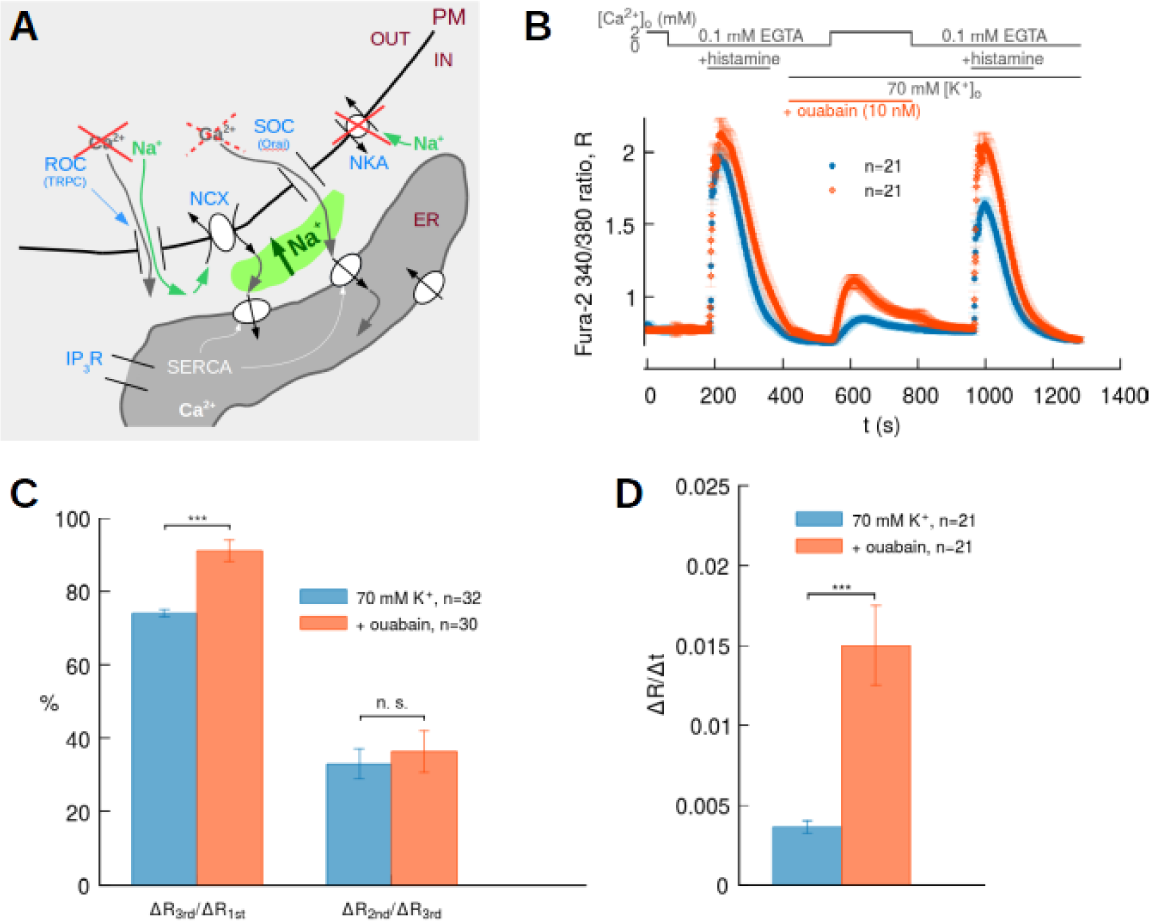
**A**. Schematics of the possible ion fluxes at PM-ER junctions during ouabain inhibition of low ouabain affinity NKA*α*_2_ and NKA*α*_3_ pumps. **B**. Effect of 10 nM ouabain on ER refilling (open orange diamonds); recordings from our standard protocol (solid blue circles) reported as reference; **C**. Bar chart from traces in B; left, ratio between 3^rd^ and 1^st^ Ca^2+^ transient amplitudes in A; right, ratio between 2^nd^ and 3^rd^ Ca^2+^ transient amplitudes in B; **D**. Bar chart from traces in B, comparing the Ca^2+^ re-entry rates at the start of the nCa^2+^ phase of the protocol in A.

### 3.4 SOCE (Orai1 channels) and NCX joint and sole contributors to ER Ca^2+^ refilling

It is evident from the results reported in figure 3 that a substantial fraction of ER Ca^2+^ is still releasable from the ER even after inhibition of NCX Ca^2+^ entry. This is likely only due to SOCE, since under membrane depolarization conditions and in absence of cell stimulation (note that histamine washout begins 3 min before nCa^2+^ phase), any contribution to Ca^2+^ entry by the TRPC should be negligible. Therefore, in spite of earlier indications that membrane depolarization by high [K^+^]_o_should also substantially reduce or even eliminate Ca^2+^ entry by Orai1 channels, we performed a series of NCX Ca^2+^ entry mode inhibition experiments on a population of cells, which were deprived of functional Orai1 channels by means of transfection with Orai1^dn^. This experimental condition should ensure inhibition and elimination of NCX and SOCE extracellular Ca^2+^ entry pathways, respectively. Results obtained with this protocol are reported in figure 5. The nCa^2+^ phase of the mean trace in figure 5B resembles closely the findings from the protocol in which the nCa^2+^ was omitted (red data in figure 2). The amplitude ratio between the 2^nd^ and 1^st^ histamine transients is further attenuated compared to that of the NCX inhibition experiments. The indication from these experiments is therefore that Orai1 and NCX contribute the vast majority of the ER Ca^2+^ refilling in depolarized EC.

**Figure 5:**
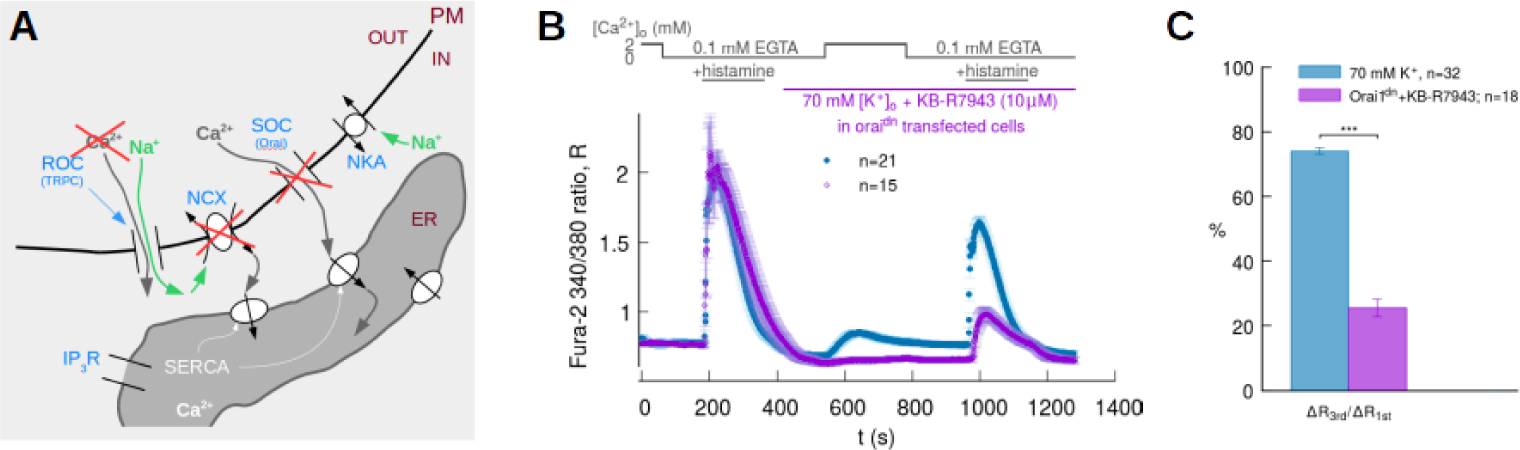
**A**. Schematics of the possible ion fluxes in PM-ER junctional regions during NCX inhibition in Orai1^dn^-transfected cells. **B**. Effect of 10 *µ*M KB-R7943 on ER refilling in absence of SOCE contribution by the STIM-Orai1 system (open purple diamonds); recordings from our standard protocol (solid blue circles) reported as reference; **C**. Bar chart from traces in B.

### 3.5 Ultrastructural evidence of PM-ER junctions in EA.hy926 cells

Since the collection of functional observations on Ca^2+^ handling to this point strongly suggests that the ER Ca^2+^ refilling process is assisted by PM-ER junctions, we surveyed a set of 29 electron micrographs, like the one reported in figure 6A, from 12 different EA.hy926 cells, to verify that PM-ER junctions are observable in the peripheral architecture of these cells.

Image analysis revealed 41 instances of junctions with mean PM-ER separation and linear apposition extension of about 10 nm and 130 nm, respectively. Moreover, in at least 60% of the junctions we observed electron opaque pillar-like structures spanning the gap of the junction and somewhat irregularly spaced, akin to those reported in other examples of S/ER junctions [38, 39] (figure 6 and table 1). These features may in reality be present more frequently than we were able to quantify, since in a number of micrographs, although the presence of a junction is clear, the PM results too smeared to establish whether pillars are present or not. (The smearing effect is probably due to the fact that cultured cells lie flat on the sample substrate and present a cross-section that is bulbous at the location of the nucleus and gets progressively thinner toward the cell periphery. In these conditions, when sectioning the samples in a direction parallel to the cell substrate it becomes less likely that the knife cuts the PM as cleanly as other intracellular membranes lying in the thicker part of the cells. For this reason, we also opted to re-orient the samples so that they could be sectioned in a direction perpendicular to the substrate plane.)

**Figure 6:**
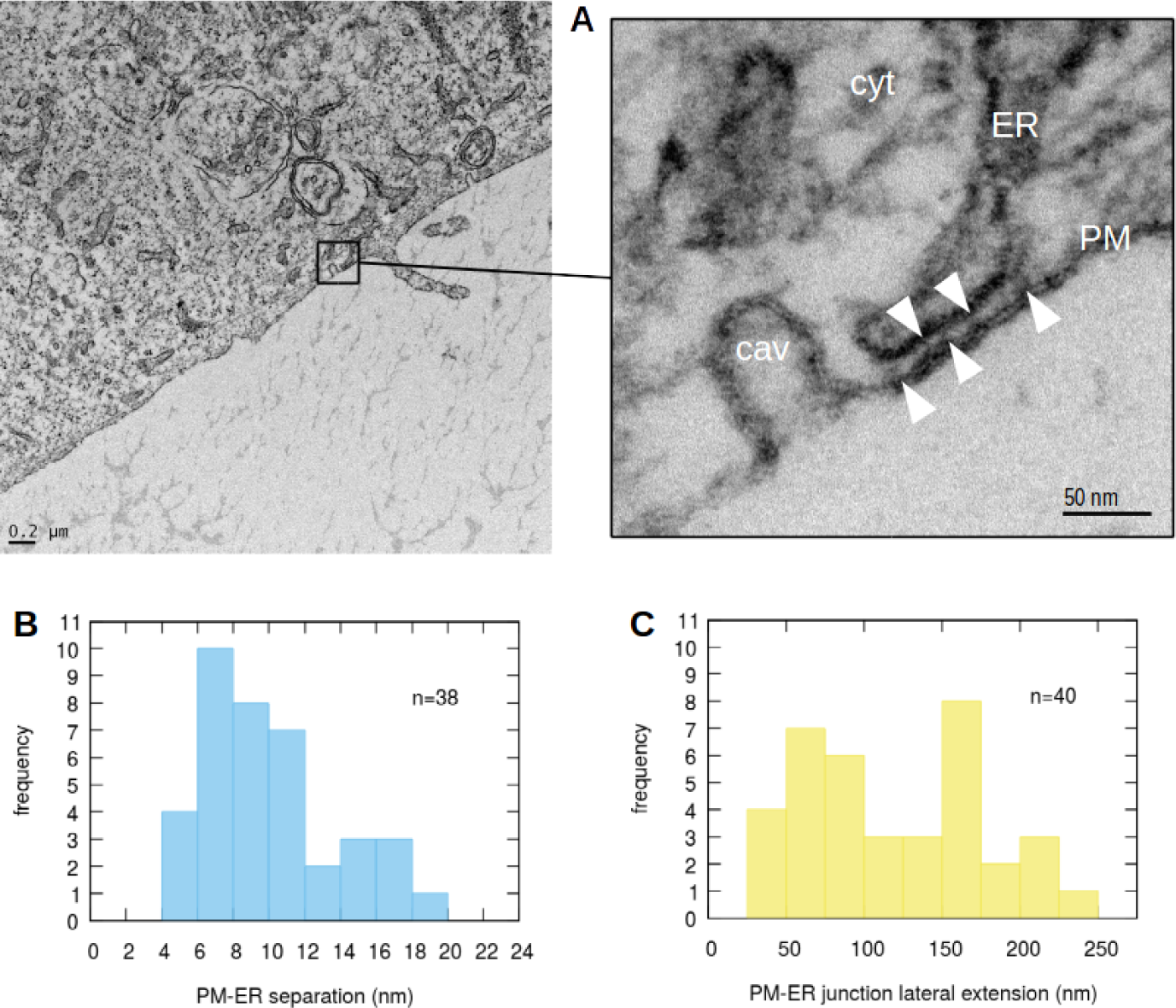
**A**. Representative electron micrograph of a peripheral region of an EA.hy926 cultured endothelial cell; the right side panel is a higher resolution image of part of the same region showing a PM-ER junction near a caveola and several instances of junction-spanning electron opaque structures (white arrowheads; cyt = cytoplasm, cav = caveola); **B. and C.** Histograms showing the distributions of PM-ER widths and lateral extensions, respectively.

**Table 1.** Statistics on ultrastructural image analysis

|  | PM-ER<br>separation (nm) | PM-ER linear<br>lateral extension (nm) | pillar<br>thickness (nm) | pillar<br>separation (nm) |
| --- | --- | --- | --- | --- |
| **mean ± sd** or<br>**observed range** | $9.6 \pm 3.8$ | $128 \pm 63$ | $4.8 \pm 1.3$ | 9–58 |

## 4 Discussion

The results reported in this article demonstrate that, following IP_3_R-mediated ER Ca^2+^ release, Ca^2+^ refilling occurs chiefly by Ca^2+^ entering the cytoplasm concomitantly via the NCX in Ca^2+^ entry mode and through the SOCE Orai1 channels.

Importantly, our findings also provide a clear indication that the refilling process takes Ca^2+^ from the extracellular space to the ER via PM-ER junctions with the NCX and Orai1 sharing the transport duties almost equally.

In line with this concept of junctional ER refilling, earlier work done under slightly different conditions suggested an ER refilling mechanism in this cell type acting in a manner that was undetectable in the bulk Fura-2 signal [5]. The specific Ca^2+^ transport players were not identified at that time, other than suggesting ER refilling was due to SOCE.

Using an external buffer containing 70 mM K^+^, we reduced the electrochemical gradient to a level expected to eliminate Ca^2+^ entry via the non-selective TRPC channels. Our main protocol in these conditions allowed us to single out the contributions of the NCX and Orai1 to Ca^2+^ entry during the nCa^2+^ phase (figure 2C and D, solid blue circles). A comparison between data from this protocol and those obtained with a physiological superfusate containing 5 mM K^+^ (figure 2B, C and D, solid black circles) strongly suggests that the ER refilling mechanism is still fully functional, even after Ca^2+^ entry pathways related to TRPC and similar non-selective cation channels are suppressed. This is revealed by the not significantly different magnitude of the second ER Ca^2+^ release transient amplitude between the normally polarized and depolarized protocols (black and blue bars in figure 2D). Furthermore, since the Fura-2 signals we report are a global measure of [Ca^2+^]_i_ in the cell, these results indicate that the refilling must take place in a manner that all but by-passes the bulk cytosol. One way for this to occur is via PM-ER nanojunctions, in which, due to their sheer dimensions, Ca^2+^ transients are undetectable by any currently available Ca^2+^ imaging technique.

Following our process of elimination of possible sources of ER Ca^2+^ refilling, in another set of experiments, we inhibited the Ca^2+^ influx mode of the NCX with a low dose of KB-R7943 (10 *µ*M [40]), still under depolarizing conditions. The most prominent feature of the observed signal is a markedly reduced amplitude of the second ER Ca^2+^ release transient (figure 3B and 3C left) showing that inhibiting Ca^2+^ entry via NCX removes a Ca^2+^ source which contributes about 40% of the ER releasable Ca^2+^ content. Moreover, we notice no significant difference between the magnitude ratios of the nCa^2+^ phase to the 2^nd^ ER release transients in the experiments with and without NCX inhibition (figure 3C right), but a significant difference between the rates of Ca^2+^ entry at the beginning of the nCa^2+^ phase of the protocol (figure 3D). Therefore, while substantial for the refilling of the ER, the effect of NCX inhibition on the bulk cytosolic signal is merely to slow down cytosolic Ca^2+^ re-entry without much altering the ratio between the accumulation in the bulk and that in the ER.

To further characterize the role of the NCX, we created a condition to amplify its Ca^2+^ influx mode by blocking the low affinity NKA*α*_2_ and NKA*α*_3_ pumps with 10 nM ouabain. If, like in vascular smooth muscle, these isoforms were primarily distributed in the PM-ER junctional regions of vascular endothelial cells [35–37], this choice would cause a strong Na^+^ transient in the junctions provoking, in turn, an increased Ca^2+^ entry through the NCX and into the ER (figure 4A). This is indeed what we observed as reported in figure 4 (B and C left), that is, a significantly larger amplitude ratio between the 2^nd^ and 1^st^ histamine-stimulated ER Ca^2+^ release transients. However, also in this experimental situation, we find that the nCa^2+^ phase transient relationship to the ER Ca^2+^ releasable fraction is not significantly different from the one observed in the control set (figure 4 C right), while the rate of Ca^2+^ entry during the nCa^2+^ phase is significantly steeper (figure 4 D).

Taken together the findings from the NCX Ca^2+^ influx mode inhibition and amplification experiments are consistent with a picture suggesting that the extracellular Ca^2+^ entering through the NCX is for the most part directly sequestered in the ER, with perhaps a minor amount contributing to the cytosolic [Ca^2+^]. Moreover, the high-ouabain-affinity-NKA inhibition experiments also support the conjecture that the NCX Ca^2+^ transport takes place at PM-ER junctions. In this view, the fraction of bulk [Ca^2+^]_i_ contributed by the NCX appears marginal and due to spill-over from the junctions, which would agree with the observed decreased or increased Ca^2+^ re-entry rate upon NCX Ca^2+^ entry mode inhibition or amplification, respectively. In the former case, diminished spill-over would hinder Ca^2+^ accumulation rate in the bulk signal, in the latter, somewhat “overdriving” NCX-dependent Ca^2+^ influx would also increase Ca^2+^ escape from the junctions, which can manifest itself as a higher rate of entry in the cytosolic signal during the initial nCa^2+^ phase of the protocol.

When we inhibited NCX Ca^2+^ entry in a population of cells transfected with Orai1^dn^, we clearly observed that SOCE via Orai1 channels contributes almost all of the releasable fraction of ER Ca^2+^ remaining after NCX inhibition in wild type cells. This can be seen in figure 5C, where the ratio between the 2^nd^ and 1^st^ histamine-stimulated transient amplitudes is further reduced to about 30% of the control experiments in (depolarized) wild type cells. Since Orai1 Ca^2+^ transport is “by definition” junctional, Orai1 SOCE must contribute to the bulk [Ca^2+^]_i_ via spill-over from PM-ER junctions, besides partially refilling ER Ca^2+^.

These experiments also indicate that in virtual absence of non-selective cation channels (e. g., TRPC) contribution to the [Ca^2+^]_i_ signal, Orai1 channels are responsible for the near totality of the Ca^2+^ re-entry signal, which can be inferred from the complete elimination of the nCa^2+^-phase signal (figure 5B). Of note, the significant contribution of Orai1-mediated Ca^2+^ entry to junctional ER-refilling occurs even in depolarized cells with strongly reduced driving force for Ca^2+^ entry. Thus, the privileged refilling of the ER within junctional areas is likely to require minute Ca^2+^fluxes through the highly selective Orai1 channels. That TRPC channels contribute the remainder of the bulk signal observed during the nCa^2+^ phase and virtually none of the ER Ca^2+^ releasable fraction was confirmed by NCX Ca^2+^ influx mode inhibition experiments using normally polarized cells (data not shown).

Analyzing the dependence of the NCX reversal potential, *E*_NCX_, on [Ca^2+^]_i_ and [Na^+^]_i_ lets us infer that the ER refilling mechanism may be assisted by PM-ER junctions also in normally polarized cells. In figure 7 we plotted *E*_NCX_ = 3 *E*_Na_ *−* 2 *E*_Ca_, where *E*_Na_ and *E*_Ca_ are the Nernst potentials for Na^+^ and Ca^2+^, respectively, as a function of [Ca^2+^]_i_ at 3 different possible values of [Na^+^]_i_. Two values of the membrane potential of our cells are also plotted: *V*_m_ *≈ −*2 mV (blue line) is the one we expect in depolarized conditions (as determined in [28]), while *V*_m_ *≈ −*50 mV (green line) is the one calculated by Goldman’s equation, using permeability values found in the literature [41]. (Other reported *V*_m_ measurements in EC in the literature under physiological conditions indicate values virtually always around, or less negative than, *−*40 mV [33, 34, 42–46].)

Since the NCX is in Ca^2+^ influx mode when *E*_NCX_ < *V*_m_, it is evident from the plots in figure 7 that in our experimental depolarizing condition of 70 mM K^+^ (and [Na^+^]_o_ = 73 mM) and using typical bulk ionic concentrations values of [Na^+^]_i_ = 10 mM and [Ca^2+^]_o_ = 2 mM, the NCX can be expected always to function in Ca^2+^ entry mode at bulk [Ca^2+^]_i_ = 100 nM (the curve corresponding to *E*_NCX_ calculated with [Na^+^]_i_ = 10 mM is well below the depolarized *V*_m_ at [Ca^2+^]_i_ = 100 nM).

**Figure 7:**
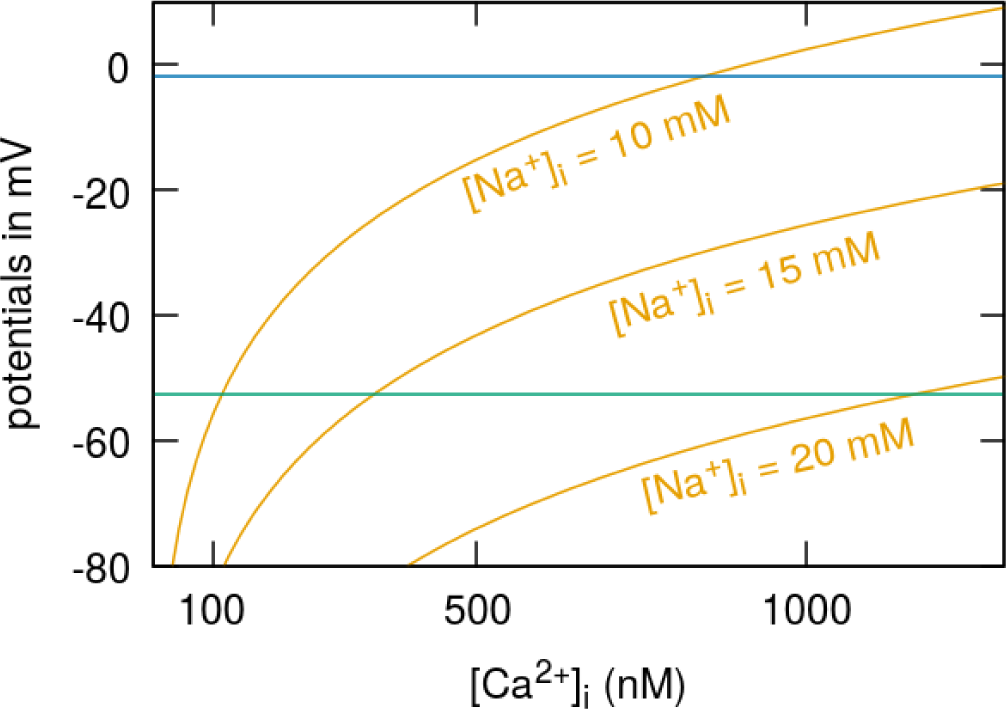
NCX reversal potential *vs* [Ca^2+^]_i_, *V*_m_ as measured in [28] (blue line) and calculated by the Goldman’s equation (green).

On the other hand, under physiological conditions of bulk cytoplasmic and of extracellular ionic concentrations normally observed in endothelial cells, the *V*_m_ being much more negative implies that the NCX reversal condition is barely respected at bulk [Ca^2+^]_i_ = 100 nM and [Na^+^]_i_ = 10 mM (the [Na^+^]_i_ = 10 mM curve crosses the *V*_m_ *≈ −*50 mV line in the neighbourhood of [Ca^2+^]_i_ = 100 nM). As the plots suggest, assuming that [Ca^2+^]_i_ equals its bulk value of 100 nm, the condition would be more safely obeyed at higher values of [Na^+^]_i_ (see for example the curve corresponding to *E*_NCX_ calculated with [Na^+^]_i_ = 15 mM). However, according to presently available data, we can expect to observe such high Na^+^ transients only in subplasmalemmal regions of cells, which are hypothesized to correspond to S/ER junctions [47, 48]. But [Ca^2+^]_i_ in such junctions is highly variable due to their sheer dimension and can easily reach micro-molar values [49]. In this case, the plot in figure 7 indicates that values of [Na^+^]_i_ closer to 20 mM would be necessary to achieve NCX reversal, which further confirms the conjecture arrived at by our functional measurements, in the depolarized system, that the NCX-mediated ER Ca^2+^ refilling mechanism must occur via PM-ER junctions in normally polarized EC as well. (While a useful guideline, it is to a certain extent deceiving to plot continuous curves to represent the variation of *E*_NCX_ as a function of [Ca^2+^]_i_ since the latter is not known to vary continuously within the cytosol. Rather we expect [Ca^2+^]_i_ *∼* 100 nM and stable in the bulk cytosol, but *∼* 1–10 *µ*M and highly variable in junctional environments.)

The unambiguous indication from our Fura-2 measurements, as well as from our analysis of the electrochemical potential, that ER Ca^2+^ refilling occurs via PM-ER junctions in a privileged manner partly segregated from bulk cytosolic Ca^2+^ prompted us to study the ultrastructure of EA.hy926 cells in search for occurrence of such junctions. A survey of 29 electron micrographs suggests that peripheral ER appositions to the PM occur regularly in these cells and possess features also observed in S/ER junctions of other cells systems, which are involved in privileged Ca^2+^ signaling, such as the cardiac dyads [50], the mitochondria-ER junctions (also known as mitochondrial associated membranes (MAMs)) [51], the PM-SR junctions in vascular smooth muscle [52, 53]. We found that PM-ER junctions in EA.hy926 cells are roughly 130 nm in extension and present a mean PM-to-ER separation of about 10 nm, having thereby a ratio between separation and extension of less than 1:10 (figure 6 and table 1). The further observation of junction-spanning structures in the majority of the analyzed junctions suggests, as in other cases, that these approaches of the ER to the PM are likely not random juxtapositions, but rather stable, if dynamic, features possibly able to fulfil specified functions, like the hypothesized privileged ER Ca^2+^ refilling. The architectural features of the PM-ER junctions we report herein would enable them to provide the tightly confined environment necessary to generate spatially and temporally localized high [Ca^2+^]_i_and [Na^+^]_i_ transients putatively responsible for Ca^2+^ signal segregation leading to the focal ER refilling we infer from our data. Confirmation of this hypothesized role of the PM-ER junctions in EC will only be possible by a combination of specific labeling of the NCX, Orai1 and TRPC and realistic quantitative modeling of the junctional transport dynamics.

In conclusion, our findings based on previously unavailable specific Ca^2+^ signal measurements and ultrastructural observations in the human umbilical vein-derived cell line EA.hy926 elucidate important steps in the mechanism of ER Ca^2+^ refilling and indicates that most of the ER Ca^2+^ releasable fraction is transported from the extra-cellular space to the ER lumen by the NCX and the Orai1 channels in similar proportions. Furthermore, the transport takes place via PM-ER junctions, which we identified and characterized for the first time. Our data therefore suggest that PM-ER junctions in EC can constitute an active functional unit for directed Ca^2+^ transport much like the hypothesized SCCU [20] and the superficial buffer barrier [22].

## Acknowledgements

We acknowledge the feedback and input from Gerd Leitinger, and the invaluable technical support of Elizabeth Pritz, Michaela Janschitz, Ren´e Rost and Anna Schreilechner. Lastly, we express our gratitude to C. J. Edcell for generating the initial EA.hy926 cell population.

